# Clinical epidemiology of congenital heart diseases in dogs: prevalence, popularity and volatility throughout twenty years of clinical practice

**DOI:** 10.1101/2020.02.25.964262

**Authors:** PG Brambilla, M Polli, D Pradelli, M Papa, R Rizzi, M Bagardi, C Bussadori

**Affiliations:** Department of Veterinary Medicine, University of Milan, Via dell’Università n. 6, 26900 Lodi (LO) - Italy; Department of Cardiology, Clinica Veterinaria Gran Sasso, Milan, Italy

## Abstract

The epidemiology of Congenital Heart Diseases (CHDs) has changed over the past twenty years.

We evaluated the prevalence of CHDs in the population of dogs recruited in a single referral center (RC); compared the epidemiological features of CHDs in screened breeds (Boxers) versus nonscreened (French and English Bulldogs and German Shepherds), investigated the association of breeds with the prevalence of CHDs, determined the popularity and volatility of breeds over a 20-year period; and analysed the trends of the most popular breeds in the overall population of new-born dogs registered in the Italian Kennel Club from 1st January 1997 to 31st December 2017.

This was a retrospective observational study, the cardiological database of the RC was analysed, and 1,779 clinical records fulfilled the inclusion criteria.

Descriptive statistics and frequencies regarding the most representative breeds and CHDs were generated. A logistic regression model was used to analyse the trends of the most common CHDs found in single breeds (French Bulldog, English Bulldog, Boxer, and German Shepherd), and in groups of breeds (brachycephalic breeds and the most represented large breeds). The relationships between the breed popularity and the presence of CHDs was studied.

The most common CHDs were Pulmonic Stenosis (34,1%), Patent Ductus Arteriosus (26,4%), Subaortic Stenosis (14,6%), Ventricular Septal Defect (4,8%), Aortic Stenosis (4,7%), Tricuspid Dysplasia (3,4%), Atrial Septal Defect (1,9%), Double Chamber Right ventricle (1,8%), Mitral Dysplasia (1,6%), and reverse Patent Ductus Arteriosus (0,7%). The most represented pure breeds were Boxer (19,4%), German Shepherd (9,4%), French Bulldog (6,2%), English Bulldog (4,9%), Maltese (3,7%), Newfoundland (3,1%), Rottweiler (3,1%), Golden Retriever (3,0%), Chihuahua (2,8%), Poodle (2,5%), Cavalier King Charles Spaniel (2,2%), American Staffordshire Terrier (2,1%), Labrador Retriever (2,3%), Dobermann (2,1%), Miniature Pinscher (2,0%), Cocker Spaniel (2,0%), Yorkshire Terrier (1,7%), Dogue de Bordeaux (1,6%), Dachshund (1,6%), and Bull Terrier (1,5%). Chihuahuas, American Staffordshire Terriers, Border Collies, French Bulldogs, and Cavalier King Charles Spaniel were the most appreciated small and medium breeds, all of which showed a high value of volatility.

In conclusion, this study found evidence for the value of the screening program implemented in Boxers, which decreased the prevalence of Subaortic Stenosis and Pulmonic Stenosis. However, fashions and trends influence dog owners’ choices more than the worries of health problems frequently found in a breed. Effective breeding programs are needed in order to control the diffusion of CHDs without impoverishing the genetic pool; in addition, dog owners should be educated, and the breeders supported by a network of veterinary cardiology centers.

## Introduction

Congenital anomalies of the cardiovascular system are defects present at birth, and often lead to perinatal death in dogs. However, in some cases, congenital heart diseases are asymptomatic and undetected until later in life, so the percentage of dogs with congenital heart diseases that survive to adulthood to breed can be rather high. To decrease the incidence of CHDs in the dog population as a whole, the early identification of affected dogs could inform a breeding program. Furthermore, some of the most common CHDs could be successfully treated by surgical management, and an early diagnosis can help to provide a normal life expectancy compared to that of the untreated dogs [1]. Knowing the epidemiology of CHDs plays an important role in maintaining dog health and in preventing the diffusion of CHDs in the dog population.

Epidemiological studies on congenital heart disease in dogs have been conducted all over the world since the early 1960s [2,3].

The most valuable studies were performed in the USA, Australia, the UK, Switzerland, Sweden and Italy [4-9]. The main studies report different prevalence of CHDs in the affected breeds, depending on the popularity of the breed in a country in a given period of time [10-12]. In almost all studies, the most common CHDs observed were PDA, PS, and SAS [1-9]. VSD, TD and TOF have also been described by the authors of these studies, but they are not noted as frequently as the abovementioned CHDs [2,3,5,7,8].

In 2011, a retrospective epidemiological study on CHDs was performed in Italy by Oliveira et al. The included data were collected at a single veterinarian referral center for cardiovascular disease in small animals, specializing in the surgical and interventional treatment of congenital heart diseases. Since 1997, the RC, IBC and FSA have also been included in a screening program that aims to reduce the prevalence of PS and SAS in Boxers, such that many breeding dogs have been screened for these conditions before breeding.

In the last seven years, new CHDs clinical cases have been reported, and this phenomenon provided a worthy opportunity to evaluate the epidemiology of CHDs in a large population of dogs in the same RC over a longer period of time.

The aims of our study were to assess the prevalence of CHDs in the population of dogs recruited in a single RC and to analyze the trends of the most popular dog breeds in the overall population of the puppies registered in the ENCI database from 1st January 1997 to 31st December 2017 [13].

The clinic’s database was updated and reanalyzed in order to investigate any changes in the epidemiological features of congenital heart diseases in non-screened (French Bulldog, English Bulldog and German Shepherd) and screened (Boxer) dogs, to determine the association of the breed with CHD and to study the popularity and volatility of the breeds over this 20-year period.

## Materials and Methods

The medical records of dogs referred for congenital heart disorders to Clinica Veterinaria Gran Sasso between 1^st^ January 1997 to 31^st^ December 2017 were retrospectively reviewed.

The population affected by CHDs was organized in two spreadsheets (isolated and associated). Dogs affected by one CHD were included in the isolated Congenital Heart Disease group, and dogs with two or more concurrent defects were included in the associated Congenital Heart Disease group.

In this study, TOF was included in the group with isolated defects, because the pathology was considered as a unique entity.

The breeds with more than 20 dogs each and the defects diagnosed in more than 10 subjects were included. The breeds with fewer than 20 dogs and the CHD with fewer than 10 animals each were named as “others”.

Subaortic stenosis Type 1, Type 2 and Type 3 were pooled in the SAS category, whereas pulmonic stenosis type A, Type B, Type M, Type BHG, Type MHG and PS R2ACA were included in the PS group [14-23].

The crossbred dogs were considered as a single group. The unknown phenotypic features of the dogs and the missing information on age and/or weight at presentation did not allow a categorization into small, medium and large breeds.

The inclusion criteria were dogs affected by CHD with complete clinical records (signalment, history, and physical examination), including thoracic radiography and echocardiography without sedation. The angiography and postmortem examination were not executed in all cases. The diagnosis of CHD was obtained by a complete transthoracic echocardiographic examination (TTE), which was performed in all patients. TTEs were carried out using commercial ultrasound equipment with mechanical transducers ranging from 2 to 10 MHz (Caris, Esaote, Florence, Italy), and then using ultrasound machines with electronic transducers also ranging from 2 to 10 MHz (Megas Esaote, Florence, Italy; Mylab30Vet, Esaote, Florence, Italy; MyLab60, Esaote, Florence, Italy, Epiq 7 Philips S.p.A., Milan, Italy).

Two-dimensional transesophageal echocardiography (TEE 2D) was executed using an omniplane transesophageal probe (Mylab30Vet, Esaote, Florence, Italy) ranging from 3 to 8 MHz Three-dimensional TEEs were performed with an echocardiography machine equipped with an omniplane transesophageal probe x7 matrix ranging from 2 to 7 MHz (Philips IE33, S.p.A., Milan, Italy) when indicated and authorized by the owner. The exams were performed, interpreted and/or reviewed by an ECVIM board-certified cardiologist (C.B.). The patients were placed in right and left lateral recumbency, and the examinations were performed according to the American Society of Echocardiography standards and guidelines and other published recommendations [24]. Angiographic procedures were also performed and/or reviewed by an ECVIM board-certified cardiologist (C.B.) with a fluoroscopy system in cases undergoing interventional percutaneous procedures or when necessary for diagnostic purposes (Villa Sistemi Medicali S.p.A., Buccinasco (MI), Italy and Digital Fluoroscopy system Philips Veradius, Milan, Italy). Postmortem examinations were performed under the supervision of C.B.

The exclusion criteria were dogs with incomplete clinical records or that were affected by acquired heart disease. The prevalence, the popularity and the volatility of the most common breeds were evaluated by using the cohort of the puppies registered in ENCI database from January 1997 to December 2017.

## Statistical Analysis

Two datasets were available for statistical analyses, and the descriptive statistics were calculated. The Kolmogorov-Smirnov test was used to assess normality of the continuous variables. For data that were not normally distributed, median and interquartile ranges (I/Q: lower and upper quartiles) are given. The Kruskal-Wallis test was used to evaluate the significance of the association among the most common CHDs and the age at presentation. The frequency distributions of all diseases in the overall sample for sex was calculated. The χ^2^ test was used to compare the occurrence of each CHD in males and females both of purebreds and crossbreds. The combinations of the observed heart defects were analyzed as groups of two, three, four and five CHDs, except TOF, which was considered separately when detected alone.

The trend of the most common CHDs found in our population from 1997 to 2017 were evaluated in single breeds (French Bulldog, English Bulldog, Boxer, and German Shepherd) and in groups of breeds (brachycephalic breeds and the most represented large breeds). The years were pooled in 7 periods for a better description of the CHDs trend: 1997-1999, 2000-2002, 2003-2005, 2006-2008, 2009-2011, 2012-2014, and 2015-2017. For each CHD, dogs were considered ‘positive’ or ‘negative’ if they were affected or were not affected by that CHD, respectively. The risk of finding dogs with a determined CHD in a specific period and for a specific breed can be estimated by the following generalized linear model

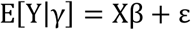

where Y is the vector of observations, β is the vector of the fixed effect (breed * period interaction) and var[ε] = var[Y| γ]. This model, applied to a binomial distribution, provides the least square means and the relative confidence intervals on a logit scale; the least square means can be reported to the probability scale by the following equation:

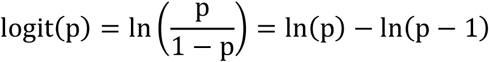

This equation can be rearranged as:

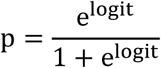

The Clinica Gran Sasso internal CHDs database was merged with the ENCI database in order to estimate the odds ratios of the overall CHD. Contingency tables were constructed for the relationships between overall CHD and each breed. The magnitude of the relationship was expressed as the odds ratio and relative 95% CI with associated P-value.

To investigate the association of CHD and the breeds’ popularity, the Pearson correlation coefficients were estimated for the total number of CHDs detected in each breed, the OR of overall CHD were determined, and two measures of breed popularity were calculated as reported by Ghirlanda et al. [10]:

1. Total popularity, defined as the total number of registrations for each breed in 1997-2017:

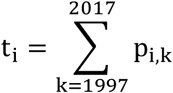
2. Volatility, defined as the average relative change in registrations from one year to the next:

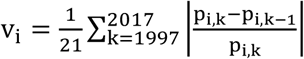

where:

t_i_ = popularity of the i^th^ breed;

v_i_ = volatility of the i^th^ breed;

p_i,k_ = number of dogs of the i^th^ breed registered in IKC in a year k (1997≤k≤2017)

21 = the number of registration changes in the period 1997-2017

For all analyses, statistical significance was set to the 5% level.

Statistical analyses were performed using the GLIMMIX, FREQ, MEANS and UNIVARIATE SAS® procedures (SAS Institute Inc. Base SAS^®^ 9.4 Procedures Guide: Statistical Procedures, Second Edition. Cary, NC: SAS Institute Inc. 2013).

## Results and Discussion

This retrospective study was based on the 1,779 clinical records that fulfilled the inclusion criteria. Single cardiac defects were present in 1,568 dogs (88.14%), and 2 or more concurrent defects were found in 211 dogs (11.86%). The total observed cases of congenital heart defects are reported in Table 1, including information on sex and age at presentation.

**Table 1.** Congenital heart defects.

| CHDs | TOTAL |  | ISOLATED |  | ASSOCIATED |  | MALE |  |  |  | FEMALE |  |  |  | AGE (months) |
| --- | --- | --- | --- | --- | --- | --- | --- | --- | --- | --- | --- | --- | --- | --- | --- |
|  | N | % | N | % | N | % | ISOLATED | ASSOCIATED | TOTAL | % | ISOLATED | ASSOCIATED | TOTAL | % | Q<sub>2</sub> (Q<sub>1</sub> – Q<sub>3</sub>)<br>9 |
| PS | 689 | 34.1 | 570 | 82.7 | 119 | 17.3 | 339 | 67 | 406 | 58.9 | 231 | 52 | 283 | 41.1 | 10 (5 - 24) |
| PDA | 534 | 26.4 | 490 | 71.1 | 44 | 8.2 | 156 | 15 | 171 | 32.0 | 334 | 29 | 363 | 68.0 | 7 (3 - 20.5) |
| SAS | 296 | 14.6 | 220 | 31.9 | 76 | 25.7 | 139 | 44 | 183 | 61.8 | 81 | 32 | 113 | 38.2 | 12.5 (4 - 31) |
| VSD | 98 | 4.8 | 39 | 5.7 | 59 | 60.2 | 21 | 33 | 54 | 55.1 | 18 | 26 | 44 | 44.9 | 8 (4 - 20.5) |
| AS | 95 | 4.7 | 80 | 11.6 | 15 | 15.8 | 54 | 7 | 61 | 64.2 | 26 | 8 | 34 | 35.8 | 25 (10 - 81) |
| TD | 69 | 3.4 | 51 | 7.4 | 18 | 26.1 | 26 | 8 | 34 | 49.3 | 25 | 10 | 35 | 50.7 | 10 (6 - 28) |
| ASD | 42 | 2.1 | 21 | 3.0 | 21 | 50.0 | 10 | 3 | 13 | 31.0 | 11 | 18 | 29 | 69.0 | 15 (7 – 35) |
| DCRV | 37 | 1.8 | 21 | 3.0 | 16 | 43.2 | 14 | 9 | 23 | 62.2 | 7 | 7 | 14 | 37.8 | 6 (4 - 11) |
| MD | 32 | 1.6 | 27 | 3.9 | 5 | 15.6 | 15 | 5 | 20 | 62.5 | 12 | 0 | 12 | 37.5 | 8 (4 - 17.5) |
| TOF | 21 | 1.0 | 0 | 0.0 | 21 | 100.0 | 0 | 11 | 11 | 52.4 | 0 | 10 | 10 | 47.6 | 5 (3 - 9) |
| rPDA | 15 | 0.7 | 15 | 2.2 | 0 | 0.0 | 6 | 0 | 6 | 40.0 | 9 | 0 | 9 | 60.0 | 11 (5 - 29) |
| MVS | 12 | 0.6 | 6 | 0.9 | 6 | 50.0 | 4 | 4 | 8 | 66.7 | 2 | 2 | 4 | 33.3 | 26.5 (3 - 76) |
| BAV | 10 | 0.5 | 5 | 0.7 | 5 | 50.0 | 3 | 5 | 8 | 80.0 | 2 | 0 | 2 | 20.0 | 14 (2 - 26) |
| PLCVC | 10 | 0.5 | 0 | 0.0 | 10 | 100.0 | 0 | 6 | 6 | 60.0 | 0 | 4 | 4 | 40.0 | 6.5 (4 - 21) |
| AVCD | 10 | 0.5 | 5 | 0.7 | 5 | 50.0 | 0 | 1 | 1 | 10.0 | 5 | 4 | 9 | 90.0 | 14 (10 - 47) |
| others<sup>a</sup> | 52 | 2.6 | 18 | 2.6 | 34 | 65.4 | 6 | 14 | 20 | 38.5 | 12 | 20 | 32 |  | 10 (4 – 54) |
|  | 2022 | 100.0 | 1568 |  | 454 |  | 793 | 232 | 1025 |  | 775 | 222 | 997 |  |  |

Legend: CHDs, congenital heart diseases; PS, pulmonic stenosis; PDA, patent ductus arteriosus; SAS, sub-aortic
stenosis; VSD, ventricular septal defect; AS, aortic stenosis; TD, tricuspid dysplasia; ASD, atrial septal defect;
DCRV, double chamber right ventricle; MD, mitral valve dysplasia; TOF, Tetralogy of Fallot; rPDA, reverse
patent ductus arteriosus; MVS, mitral valve stenosis; BAV, aortic bicuspid valve; PLCVC, persistent left cranial
vena cava; AVCD, atrioventricular canal disease;
others<sup>a</sup> CTD, cor triatriatum dexter; AVF, aortovenous fistula; PPDH, pericardial peritoneum diaphragmatic
hernia; QAV, quadricuspid aortic valve; AH, aortic hypoplasia; AI, aortic insufficiency; AOr, aortic overriding;
APW, aortopulmonary window; MI, mitral insufficiency; TVS, tricuspid stenosis; TA, truncus arteriosus; CollF, systemic to pulmonary arterial collateral flow; AC, azygos continuation; ACT, anomalous coronary truncus.

The most common CHD in our overall population, both isolated and associated with other conditions, were PS (34.1%), PDA (26.4%) SAS (14.6%), VSD (4.8%), AS (4.7%), TD (3.4%), ASD (1.9%), DCRv (1.8%), MD (1.6%), and rPDA (0.7%). Among these CHDs, the youngest dogs at presentation were affected by TOF (median 5 month) and the oldest by AS (median 25 months) (Table 1). The results from the Kruskall-Wallis test showed that the dogs diagnosed with AS were significantly older than the dogs affected by PDA, PS, AS, DCRV, VSD, SAS, TD, and TOF (Fig 1).

**Figure 1.**
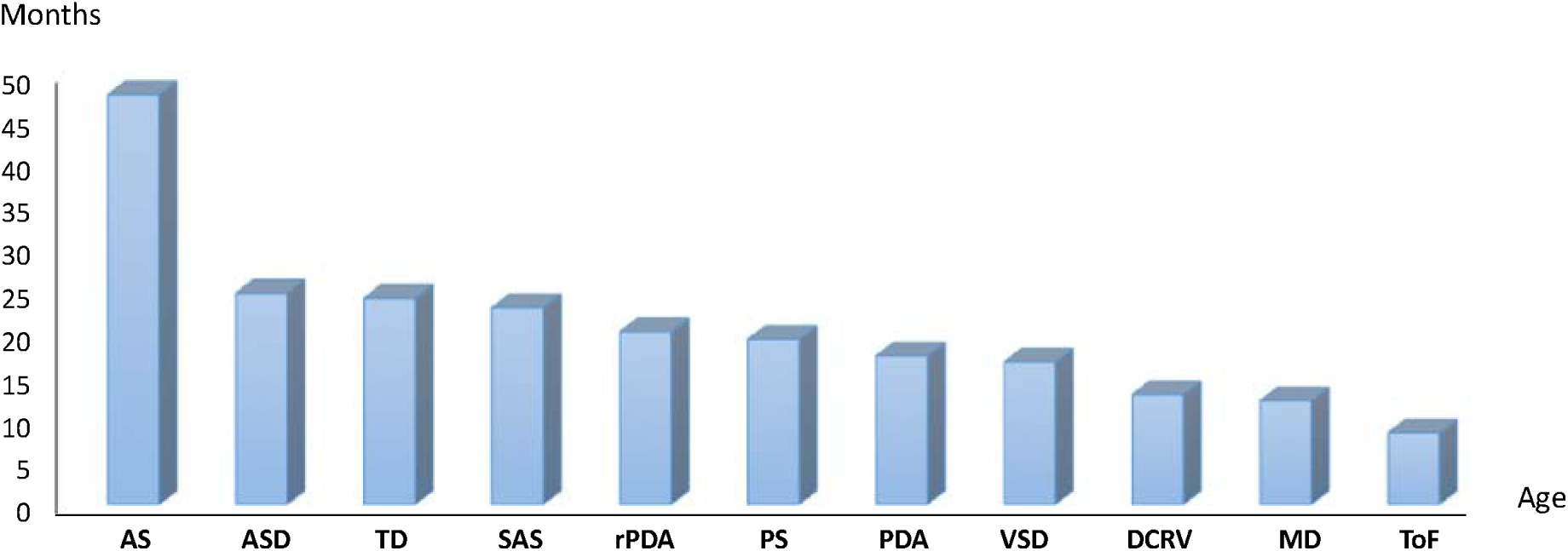
Average age (months) of the dogs belonging to the most represented isolated CHDs.

Isolated congenital heart diseases were diagnosed in 1,377 dogs belonging to 92 purebreds and 191 crossbreds. The top 21 represented purebreds were Boxer (19.4%), German Shepherd (9.4%), French Bulldog (6.2%), English Bulldog (4.9%), Maltese (3.7%), Newfoundland (3.1%), Golden Retriever (3.0%), Chihuahua (2.8%), Rottweiler (3.1%), Poodle (2.5%), Cavalier King Charles Spaniel (2.2%), American Staffordshire Terrier (2.1%), Labrador Retriever (2.3%), Dobermann (2.1%), Miniature Pinscher (2.0%), Cocker Spaniel (2.0%), Yorkshire Terrier (1.7%), Dogue de Bordeaux (1.6%), Dachshund (1.6%), and Bull Terrier (1.5%).

Of the most common CHDs found in the selected breeds, PDA was absent in Boxers, American Staffordshire Terriers and Dogue de Bordeaux. However, the same breeds experienced a large percentage of cases of PS (Boxer and American Staffordshire Terrier) and SAS (Dogue the Bordeaux) (Table 2) [25,26].

**Table 2.**
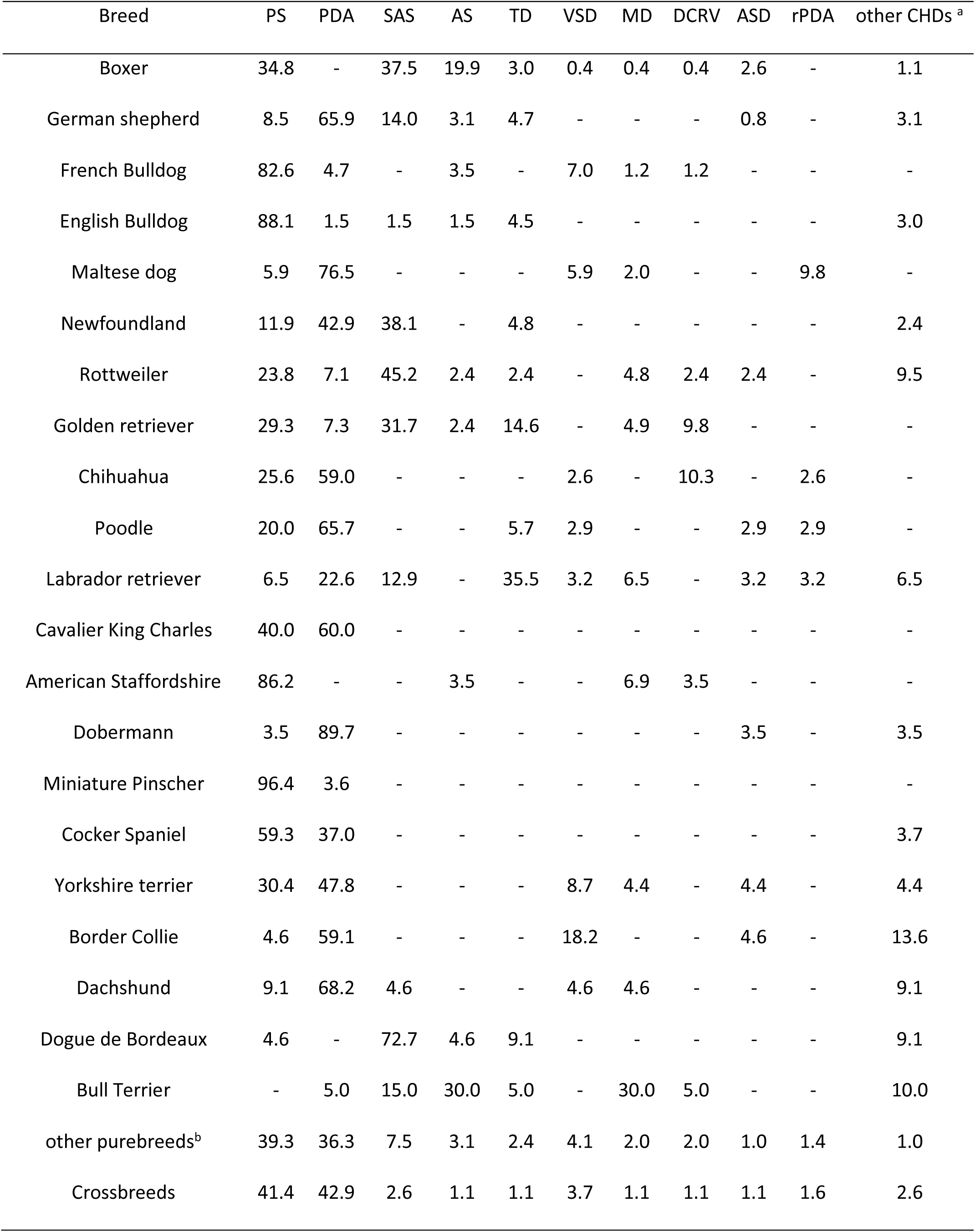
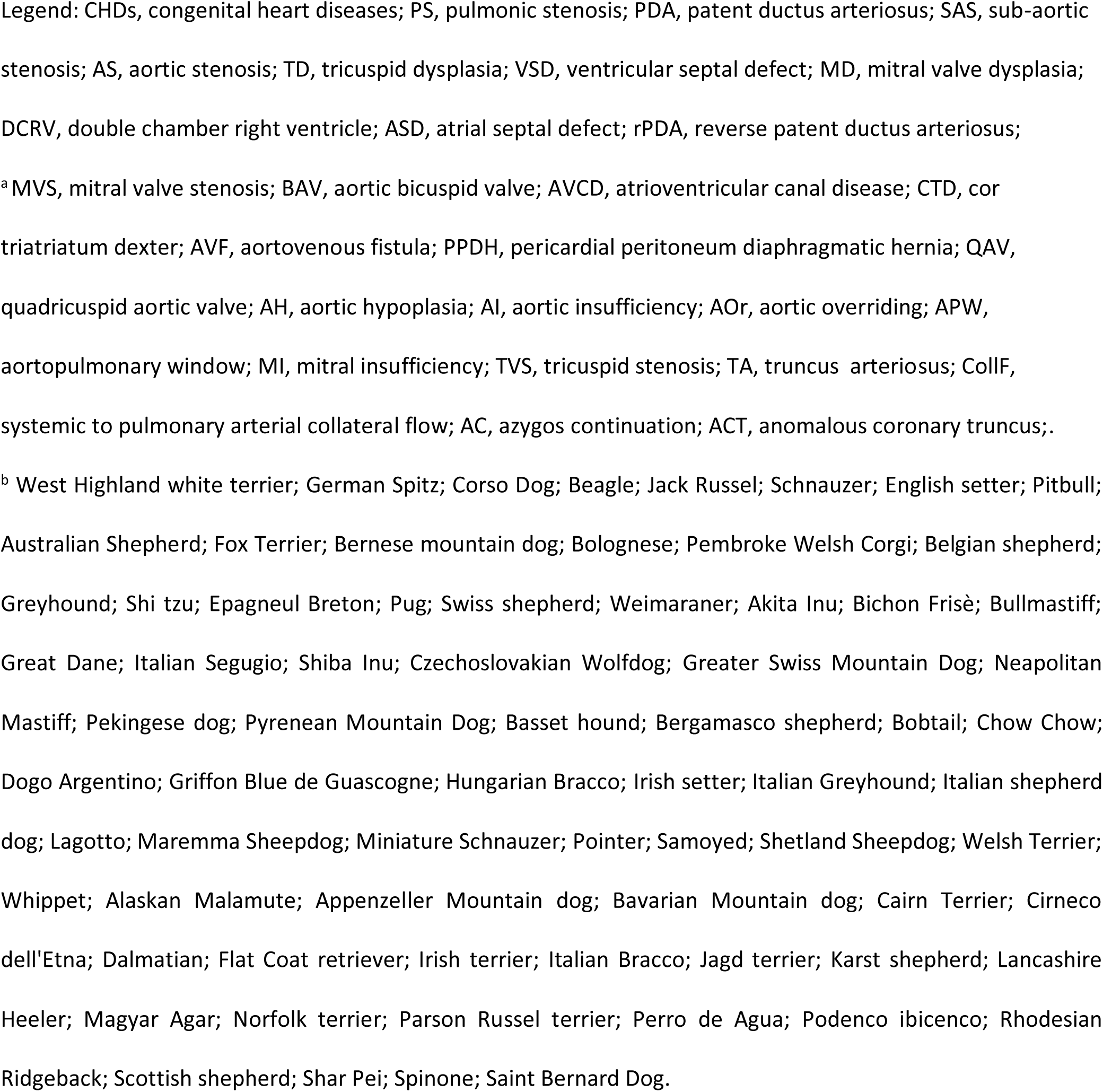
Distribution of CHDs in purebreeds and in crossbreds.

| Breed | PS | PDA | SAS | AS | TD | VSD | MD | DCRV | ASD | rPDA | other CHDs <sup>a</sup> |
| --- | --- | --- | --- | --- | --- | --- | --- | --- | --- | --- | --- |
| Boxer | 34.8 | - | 37.5 | 19.9 | 3.0 | 0.4 | 0.4 | 0.4 | 2.6 | - | 1.1 |
| German shepherd | 8.5 | 65.9 | 14.0 | 3.1 | 4.7 | - | - | - | 0.8 | - | 3.1 |
| French Bulldog | 82.6 | 4.7 | - | 3.5 | - | 7.0 | 1.2 | 1.2 | - | - | - |
| English Bulldog | 88.1 | 1.5 | 1.5 | 1.5 | 4.5 | - | - | - | - | - | 3.0 |
| Maltese dog | 5.9 | 76.5 | - | - | - | 5.9 | 2.0 | - | - | 9.8 | - |
| Newfoundland | 11.9 | 42.9 | 38.1 | - | 4.8 | - | - | - | - | - | 2.4 |
| Rottweiler | 23.8 | 7.1 | 45.2 | 2.4 | 2.4 | - | 4.8 | 2.4 | 2.4 | - | 9.5 |
| Golden retriever | 29.3 | 7.3 | 31.7 | 2.4 | 14.6 | - | 4.9 | 9.8 | - | - | - |
| Chihuahua | 25.6 | 59.0 | - | - | - | 2.6 | - | 10.3 | - | 2.6 | - |
| Poodle | 20.0 | 65.7 | - | - | 5.7 | 2.9 | - | - | 2.9 | 2.9 | - |
| Labrador retriever | 6.5 | 22.6 | 12.9 | - | 35.5 | 3.2 | 6.5 | - | 3.2 | 3.2 | 6.5 |
| Cavalier King Charles | 40.0 | 60.0 | - | - | - | - | - | - | - | - | - |
| American Staffordshire | 86.2 | - | - | 3.5 | - | - | 6.9 | 3.5 | - | - | - |
| Dobermann | 3.5 | 89.7 | - | - | - | - | - | - | 3.5 | - | 3.5 |
| Miniature Pinscher | 96.4 | 3.6 | - | - | - | - | - | - | - | - | - |
| Cocker Spaniel | 59.3 | 37.0 | - | - | - | - | - | - | - | - | 3.7 |
| Yorkshire terrier | 30.4 | 47.8 | - | - | - | 8.7 | 4.4 | - | 4.4 | - | 4.4 |
| Border Collie | 4.6 | 59.1 | - | - | - | 18.2 | - | - | 4.6 | - | 13.6 |
| Dachshund | 9.1 | 68.2 | 4.6 | - | - | 4.6 | 4.6 | - | - | - | 9.1 |
| Dogue de Bordeaux | 4.6 | - | 72.7 | 4.6 | 9.1 | - | - | - | - | - | 9.1 |
| Bull Terrier | - | 5.0 | 15.0 | 30.0 | 5.0 | - | 30.0 | 5.0 | - | - | 10.0 |
| other purebreeds<sup>b</sup> | 39.3 | 36.3 | 7.5 | 3.1 | 2.4 | 4.1 | 2.0 | 2.0 | 1.0 | 1.4 | 1.0 |
| Crossbreeds | 41.4 | 42.9 | 2.6 | 1.1 | 1.1 | 3.7 | 1.1 | 1.1 | 1.1 | 1.6 | 2.6 |

PDA was frequently found in Dobermanns, German Shepherds, Maltese and Dachshunds (Table 2). A small percentage of breeds experienced rPDA, and it was mostly detected in Maltese (9.8%) (Table 2).

In addition to Boxers, PS was very common in other brachycephalic breeds, including French and English Bulldogs. Interestingly, PS was also the most common CHD in Pinschers, in which the only other congenital heart disease observed was PDA (Table 2).

In Tables 3 and 4, the frequency of the most common CHDs by sex are reported for the purebreds and crossbreds, respectively.

**Table 3.** Distribution of CHDs by sex in purebreds.

| CHD | N | Frequency | 95% CI | Male<br>frequency | Female<br>frequency | p-value |
| --- | --- | --- | --- | --- | --- | --- |
| PS | 491 | 35.66 | 33.13 – 38.19 | 41.17 | 29.64 | <0.0001 |
| PDA | 408 | 29.63 | 27.22 – 32.04 | 19.47 | 40.73 | <0.0001 |
| SAS | 213 | 15.47 | 13.56 – 17.38 | 18.64 | 12.01 | 0.001 |
| AS | 80 | 5.81 | 4.57 – 7.05 | 7.51 | 3.95 | 0.005 |
| TD | 49 | 3.56 | 2.58 – 4.54 | 3.48 | 3.65 | 0.86 |
| VSD | 32 | 2.32 | 1.53 – 3.12 | 2.50 | 2.13 | 0.64 |
| MD | 25 | 1.82 | 1.11 – 2.52 | 1.95 | 1.67 | 0.70 |
| DCRV | 19 | 1.38 | 0.76 – 2.00 | 1.81 | 0.91 | 0.15 |
| ASD | 17 | 1.23 | 0.65 – 1.82 | 1.25 | 1.22 | 0.95 |
| rPDA | 12 | 0.87 | 0.38 – 1.36 | 0.56 | 1.22 | 0.19 |
| other CHDs <sup>a</sup> | 31 | 2.25 | 1.47 – 3.04 | 1.67 | 2.89 | 0.48 |

Legend: CHDs, congenital heart diseases; PS, pulmonic stenosis; PDA; patent ductus arteriosus; SAS, subaortic stenosis; AS, aortic stenosis; TD, tricuspid dysplasia; VSD, ventricular septal defect; MD, mitral dysplasia; DCRV, double chamber right ventricle; ASD, atrial septal defect; rPDA, reverse patent ductus arteriosus.
<sup>a</sup> MVS, mitral valve stenosis; BAV, aortic bicuspid valve; AVCD, atrioventricular canal disease; CTD, cor triatriatum dexter; AVF, aortovenous fistula; PPDH, pericardial peritoneum diaphragmatic hernia; QAV, quadricuspid aortic valve; AH, aortic hypoplasia; AI, aortic insufficiency; AOr, aortic overriding; APW, aortopulmonary window; MI, mitral insufficiency; TVS, tricuspid stenosis; TA, truncus arteriosus; CollF, systemic to pulmonary arterial collateral flow; AC, azygos continuation; ACT, anomalous coronary truncus.

**Table 4.**
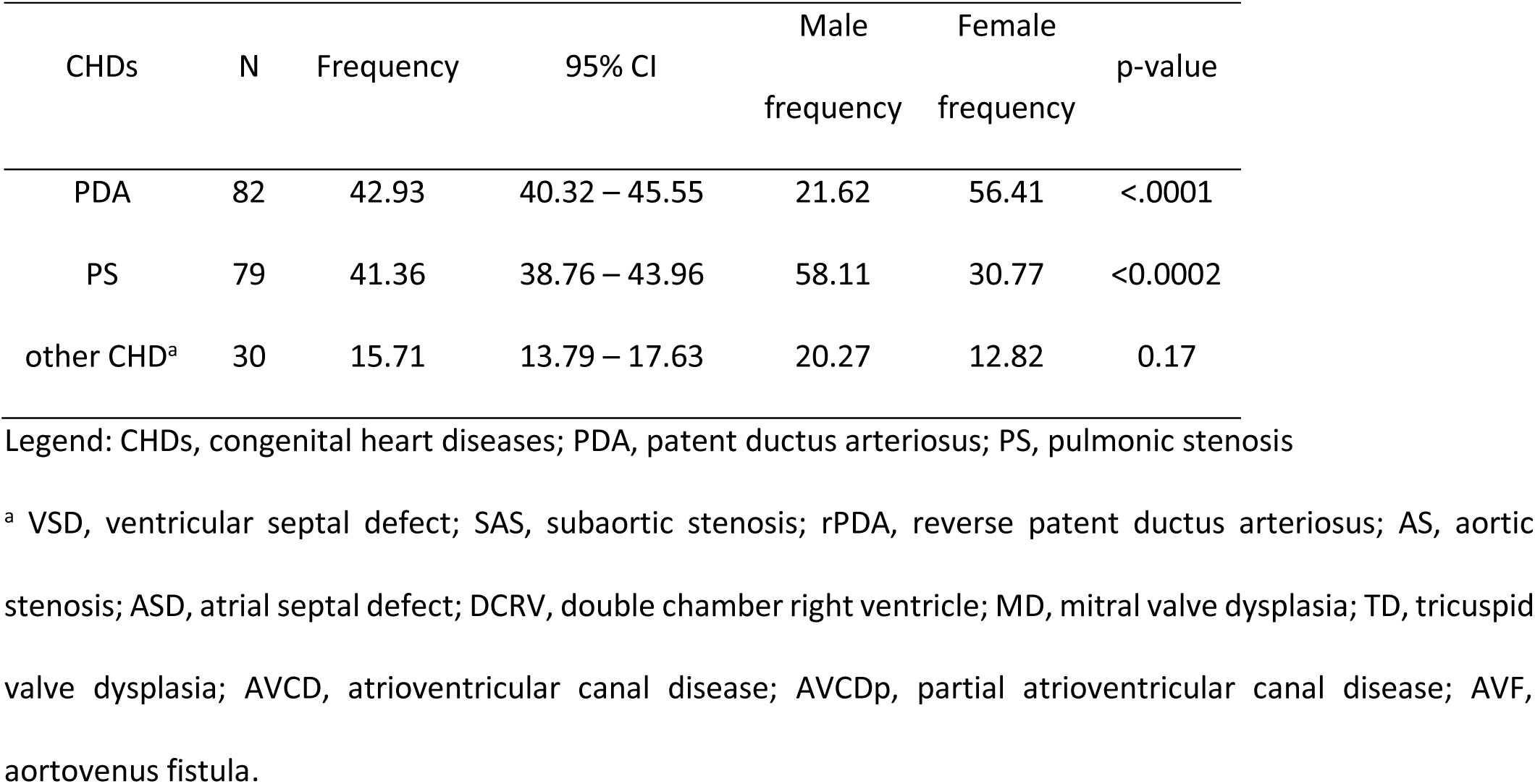
Distribution of the CHDs by sex in crossbreds.

| CHDs | N | Frequency | 95% CI | Male<br>frequency | Female<br>frequency | p-value |
| --- | --- | --- | --- | --- | --- | --- |
| PDA | 82 | 42.93 | 40.32 – 45.55 | 21.62 | 56.41 | <.0001 |
| PS | 79 | 41.36 | 38.76 – 43.96 | 58.11 | 30.77 | <0.0002 |
| other CHD<sup>a</sup> | 30 | 15.71 | 13.79 – 17.63 | 20.27 | 12.82 | 0.17 |

Legend: CHDs, congenital heart diseases; PDA, patent ductus arteriosus; PS, pulmonic stenosis
<sup>a</sup> VSD, ventricular septal defect; SAS, subaortic stenosis; rPDA, reverse patent ductus arteriosus; AS, aortic stenosis; ASD, atrial septal defect; DCRV, double chamber right ventricle; MD, mitral valve dysplasia; TD, tricuspid valve dysplasia; AVCD, atrioventricular canal disease; AVCDp, partial atrioventricular canal disease; AVF, aortovenous fistula.

In purebreds, PS, SAS and AS were significantly more frequent in males (p<0,005) while PDA was significantly more frequent in females (p<0,0001) (Table 3).

PS and PDA were also the most common cardiac defects in crossbreds, and PDA was detected significantly more frequently in females, and PS was detected more frequently in males (Table 4).

In 189 purebred dogs and in 22 crossbreds, two or more defects were detected. The most frequent association was among two simple defects (74.4%), and PS was the most frequently detected disease (59.24%). PS was associated with SAS (22.29%), VSD (19.11%) and PDA (8.28%). SAS was associated with PDA in 8.28% of the dogs (Table 5).

**Table 5.** Associations of congenital heart defects.

| CHD<sub>A</sub> |  | N.dogs | % |
| --- | --- | --- | --- |
| PS | SAS | 35 | 22.29 |
| PS | VSD | 30 | 19.11 |
| PS | PDA | 13 | 8.28 |
| PS | Other CHDs <sup>a</sup> | 15 | 9.55 |
| SAS | PDA | 13 | 8.28 |
| SAS | VSD | 5 | 3.18 |
| SAS | other CHDs <sup>b</sup> | 10 | 6.37 |
| VSD | DCRV | 7 | 4.46 |
| VSD | other CHDs <sup>c</sup> | 6 | 3.82 |
| ASD | AVCD | 5 | 3.18 |
|  | other combinations<sup>d</sup> | 18 | 11.46 |

Legend: CHD<sub>A</sub>, associated congenital heart diseases; CHD<sub>s</sub>, congenital heart diseases, PS, pulmonic stenosis; VSD, ventricular septal defect; PDA, patent ductus arteriosus;
<sup>a</sup> DCRV, double chamber right ventricle; AS, aortic stenosis; ASD, atrial septal defect; TD, tricuspid valve dysplasia; AVF, aortovenous fistula; PLCVC, persistent left atrial vena cava; TVS, tricuspid valve stenosis;
<sup>b</sup> TD, tricuspid valve dysplasia; MD, mitral valve dysplasia; MVS, mitral valve stenosis; BAV, bicuspid aortic valve; PLCVC, persistent left atrial vena cava; QAV, quadricuspid aortic valve; TOF, tetralogy of Fallot
<sup>c</sup> PDA, patent ductus arteriosus; ASD, atrial septal defect; MD, mitral valve dysplasia; AOr, aortic overriding;
<sup>d</sup> ASD-AS, atrial septal defects – aortic stenosis; ASD-TD, atrial septal defect – tricuspid dysplasia; ASD – AVCD, atrial septal defect-atrioventricular canal defects; PDA-AS, patent ductus arteriosus - aortic stenosis; PDA – ASD, patent ductus arteriosus-atrial septal defect; PDA – TD, patent ductus arteriosus-tricuspid dysplasia; PDA – MVS, patent ductus arteriosus-mitral valve stenosis; PDA – AVF, patent ductus arteriosus- aortovenous fistula; TD – DCRV, tricuspid dysplasia - double chamber right ventricle; TD – AS, tricuspid dysplasia – atrial septal defect TD – CTD, tricuspid dysplasia- cor triatriatum dexter; TOF – APW, tetralogy of Fallot-aortopulmonary window; AS – BAV, aortic stenosis-bicuspid aortic valve; DCRV - HCM; MVS – TVS, mitral valve stenosis- tricuspid valve stenosis.

There were 14.2% of dogs affected by a combination of three defects, and PS was the most frequent (61%). Eight dogs showed had four or five single defects, PS was the most frequent; it was detected in all 8 dogs.

The Tetralogy of Fallot was found in 21 (10%) dogs; 16 (7.4%) were isolated CHD conditions, and 5 (2.4%) were dogs affected by two or three single defects (Table 1).

A significant and positive relationship between the overall number of CHDs and popularity of breed was observed, suggesting that the prevalence of CHDs grows as the number of ENCI registered dogs increases (r = 0.54, p= 0.01) [13].

The number of the CHDi found in the selected breeds, the OR for the overall CHDi, the popularity of the breeds and the volatility for each breed are reported in Table 6.

**Table 6.**
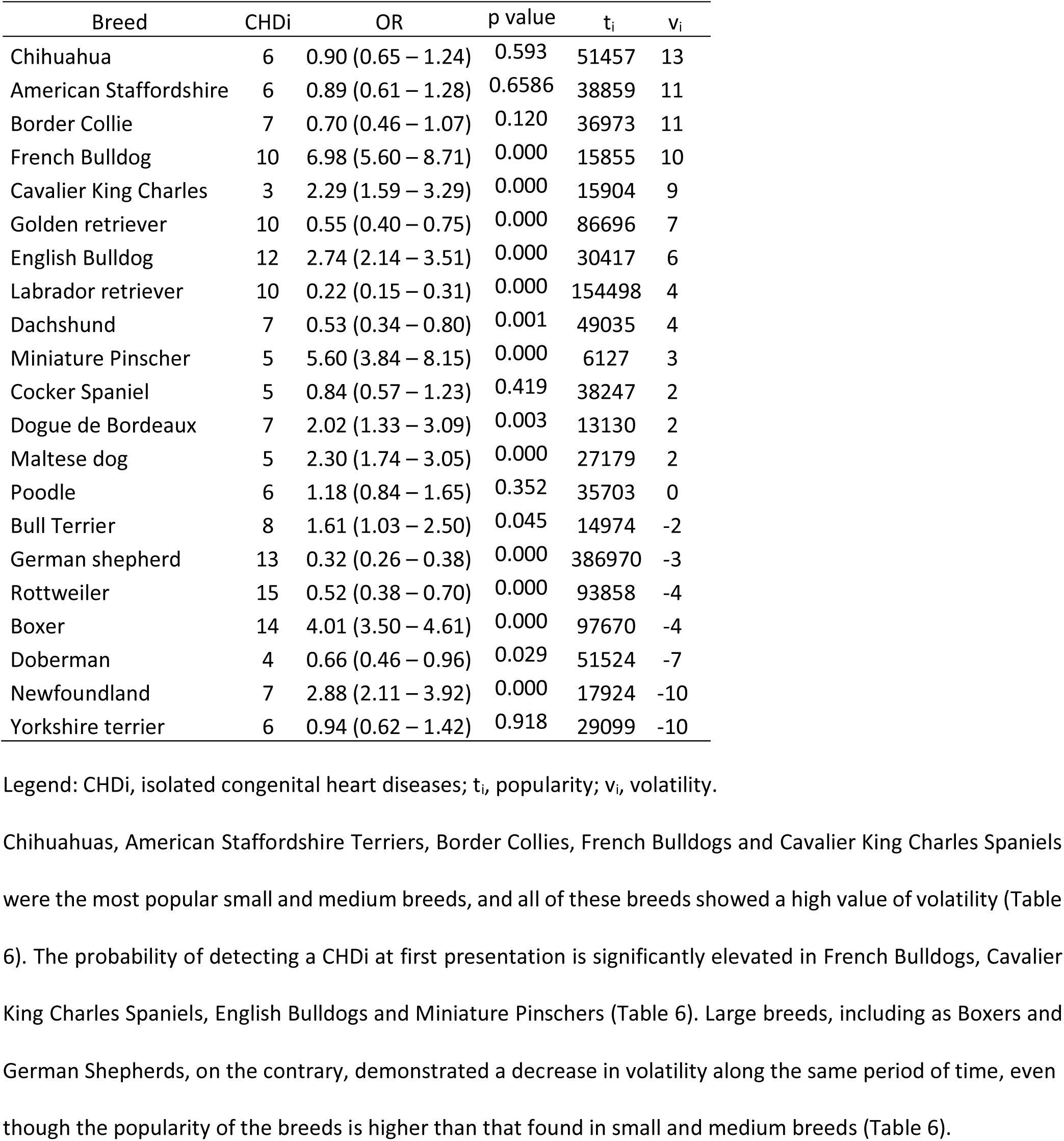
Number of CHD_i_, OR referred to overall CHD (with confidence interval), popularity, and volatility in each breed.

| Breed | CHDi | OR | p value | $t_i$ | $v_i$ |
| --- | --- | --- | --- | --- | --- |
| Chihuahua | 6 | 0.90 (0.65 – 1.24) | 0.593 | 51457 | 13 |
| American Staffordshire | 6 | 0.89 (0.61 – 1.28) | 0.6586 | 38859 | 11 |
| Border Collie | 7 | 0.70 (0.46 – 1.07) | 0.120 | 36973 | 11 |
| French Bulldog | 10 | 6.98 (5.60 – 8.71) | 0.000 | 15855 | 10 |
| Cavalier King Charles | 3 | 2.29 (1.59 – 3.29) | 0.000 | 15904 | 9 |
| Golden retriever | 10 | 0.55 (0.40 – 0.75) | 0.000 | 86696 | 7 |
| English Bulldog | 12 | 2.74 (2.14 – 3.51) | 0.000 | 30417 | 6 |
| Labrador retriever | 10 | 0.22 (0.15 – 0.31) | 0.000 | 154498 | 4 |
| Dachshund | 7 | 0.53 (0.34 – 0.80) | 0.001 | 49035 | 4 |
| Miniature Pinscher | 5 | 5.60 (3.84 – 8.15) | 0.000 | 6127 | 3 |
| Cocker Spaniel | 5 | 0.84 (0.57 – 1.23) | 0.419 | 38247 | 2 |
| Dogue de Bordeaux | 7 | 2.02 (1.33 – 3.09) | 0.003 | 13130 | 2 |
| Maltese dog | 5 | 2.30 (1.74 – 3.05) | 0.000 | 27179 | 2 |
| Poodle | 6 | 1.18 (0.84 – 1.65) | 0.352 | 35703 | 0 |
| Bull Terrier | 8 | 1.61 (1.03 – 2.50) | 0.045 | 14974 | -2 |
| German shepherd | 13 | 0.32 (0.26 – 0.38) | 0.000 | 386970 | -3 |
| Rottweiler | 15 | 0.52 (0.38 – 0.70) | 0.000 | 93858 | -4 |
| Boxer | 14 | 4.01 (3.50 – 4.61) | 0.000 | 97670 | -4 |
| Doberman | 4 | 0.66 (0.46 – 0.96) | 0.029 | 51524 | -7 |
| Newfoundland | 7 | 2.88 (2.11 – 3.92) | 0.000 | 17924 | -10 |
| Yorkshire terrier | 6 | 0.94 (0.62 – 1.42) | 0.918 | 29099 | -10 |

Legend: CHDi, isolated congenital heart diseases; $t_i$ , popularity; $v_i$ , volatility.

To the best of our knowledge, this is the longest epidemiological study on CHDs performed in dogs from a single referral center.

The clinic involved in this study is a valuable site from which to monitor the evolution of trends among different breeds because it has been a referral center for CHDs studies in the authors’ country since 1997 and is based in a large city in the northern part of the country. In accordance with other studies, PS, PDA, SAS and AS were the most common CHDs in the purebred population, PDA and PS were prevalent in crossbreds, and in both groups, males were significantly more frequently affected by CHDs than females [4-9,27]. There were, however, some differences concerning the prevalence of CHDs in the different breeds. The most common CHD in this study proved to be PS, as a single or complex defect, associated with SAS (22.29%), VSD (19.11%), PDA (8.28%) and less common CHDs (9.55%) (Tables 1 and 5). The main breeds affected by all Types of PS were brachycephalic breeds. English Bulldogs (88.1%) and French Bulldogs (82%), had the greatest prevalence of PS from the beginning of the study, and the prevalence increased over time as the Boxer PS prevalence decreased (Fig 2).

**Figure 2.**
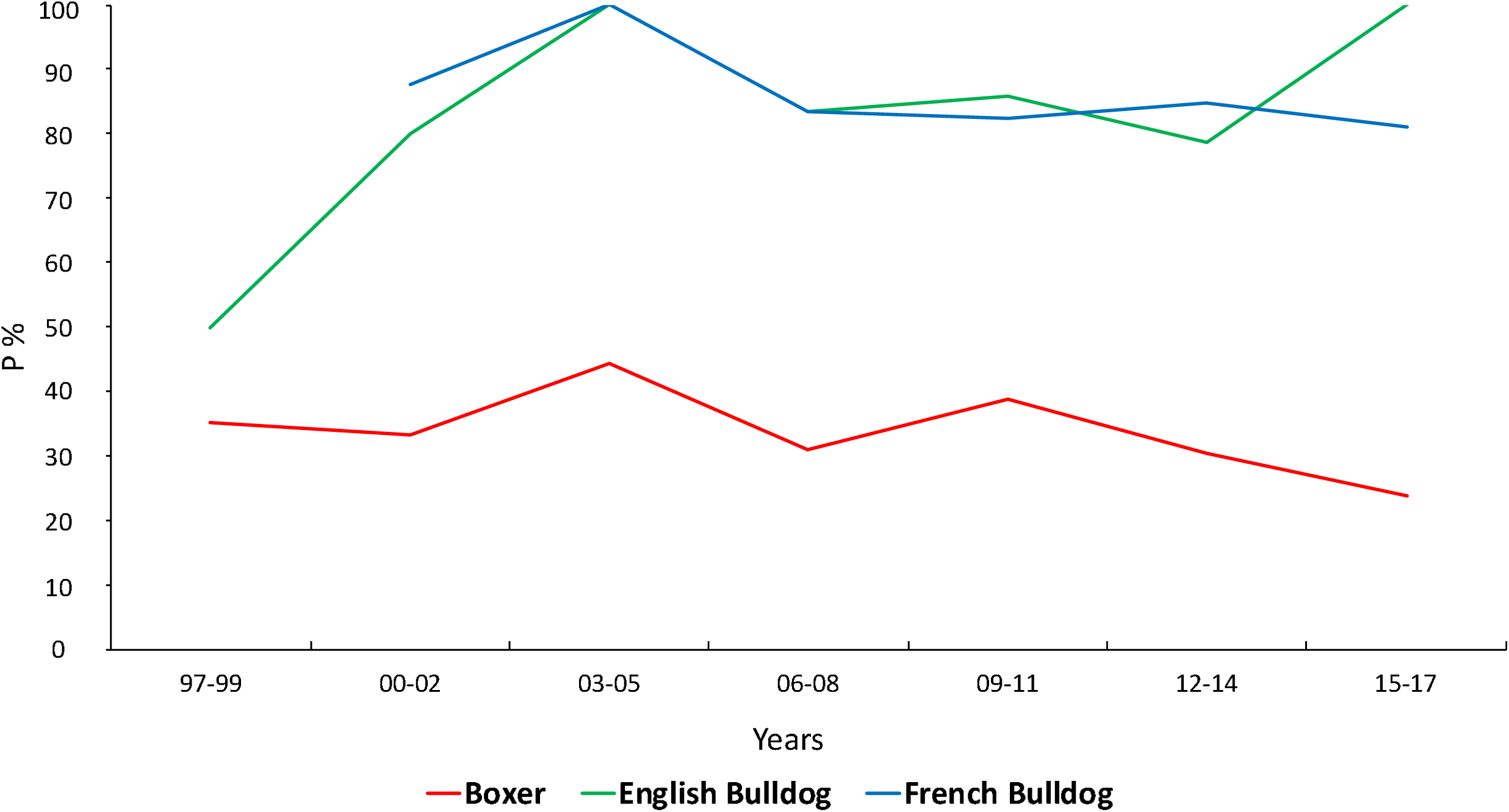
Probability of identifying Pulmonic Stenosis in Boxers, French Bulldogs and English Bulldogs admitted from 1997 to 2017. Abbreviation: Probability, P.

The most common PS Types found in the aforementioned breeds were Type A in French Bulldogs (42.25%), Type A equal to Type B in English Bulldogs (40.68%) and Type A in Boxers (59.14%).

In Boxers, only Types A and B of PS were found, while other Types were found in French Bulldogs (PS Type BHG 12.68%; PS Type M 15.49%) and in English Bulldogs (PS Type BHG, PS Type M and PS Type MHG, all together 18.64%)[15]. Although most common in English Bulldogs, PSR 2ACA was also found in French Bulldogs, Brussels Griffons, American Staffordshire Terriers and Corso dogs [20].

The probability of admitting a Boxer affected by PS decreased from 1997 (35%) to 2017 (23.8%) in the overall population of the RC (Fig 2). This result can be explained as an effect of the screening program that has been in effect since 2000 in the RC. In collaboration with BCI and FSA, the screening program collected, in a separate database, the individual phenotypic information on the traits leading to a PS diagnosis, which then gradually led to a reduction of Boxers affected by PS [18,28]. The increased number of veterinary centers qualified to perform the screening before breeding Boxers could also be a reason to explain the reduction of incoming Boxers affected by PS in the authors’ RC.

In the last decade, English Bulldogs and French Bulldogs have been dramatically increasing in popularity in our country, as observed in this study’s population. Decreased popularity of Boxers was a trend that was observed in the results published by other authors and in our clinic, as mentioned above [10,29,30].

The factors that influenced the success of brachycephalic breeds are well known by authors in UK, Denmark and the USA, where many studies have been conducted [30,31]. The lovers of the brachycephalic breeds were less influenced by health and longevity in terms of breed selection compared with non-brachycephalic dogs’ owners. A variety of different drivers have been identified to explain the popularity of English Bulldogs and French Bulldogs, including factors that influenced owners’ decisions to buy brachycephalic dogs. The breeds’ appearances (large forehead, big eyes, round face, and bulging cheeks), good behavior, deeply affectionate temperament and good relationships with children have been described as the most important determinants driving people’s desire for these breeds [10,30,31].

All of the typical brachycephalic features valued by the owners of Boxers can be found in English Bulldogs and French Bulldogs; however, the Bulldogs have a breed size more suitable to current lifestyles. The large size of Boxers could somewhat influence buyers’ breed choice and may explain the decline of this breed in our clinical setting.

PS in American Staffordshire Terriers (86.2%) and Golden Retrievers (29.3%) progressively increased from 1997 to 2017; this observation is in contrast with our explanation for the decrease in Boxer popularity regarding their size. In fact, even if the breeds are medium or large in size, they are very different from Boxers, not interchangeable, and their success was because they became fashionable. For example, American Staffordshire Terriers, are a status symbol among some young people groups’ in large European cities, and the success of Golden Retrievers was due more to the influence of movies (Fig 3) [12].

**Figure 3.**
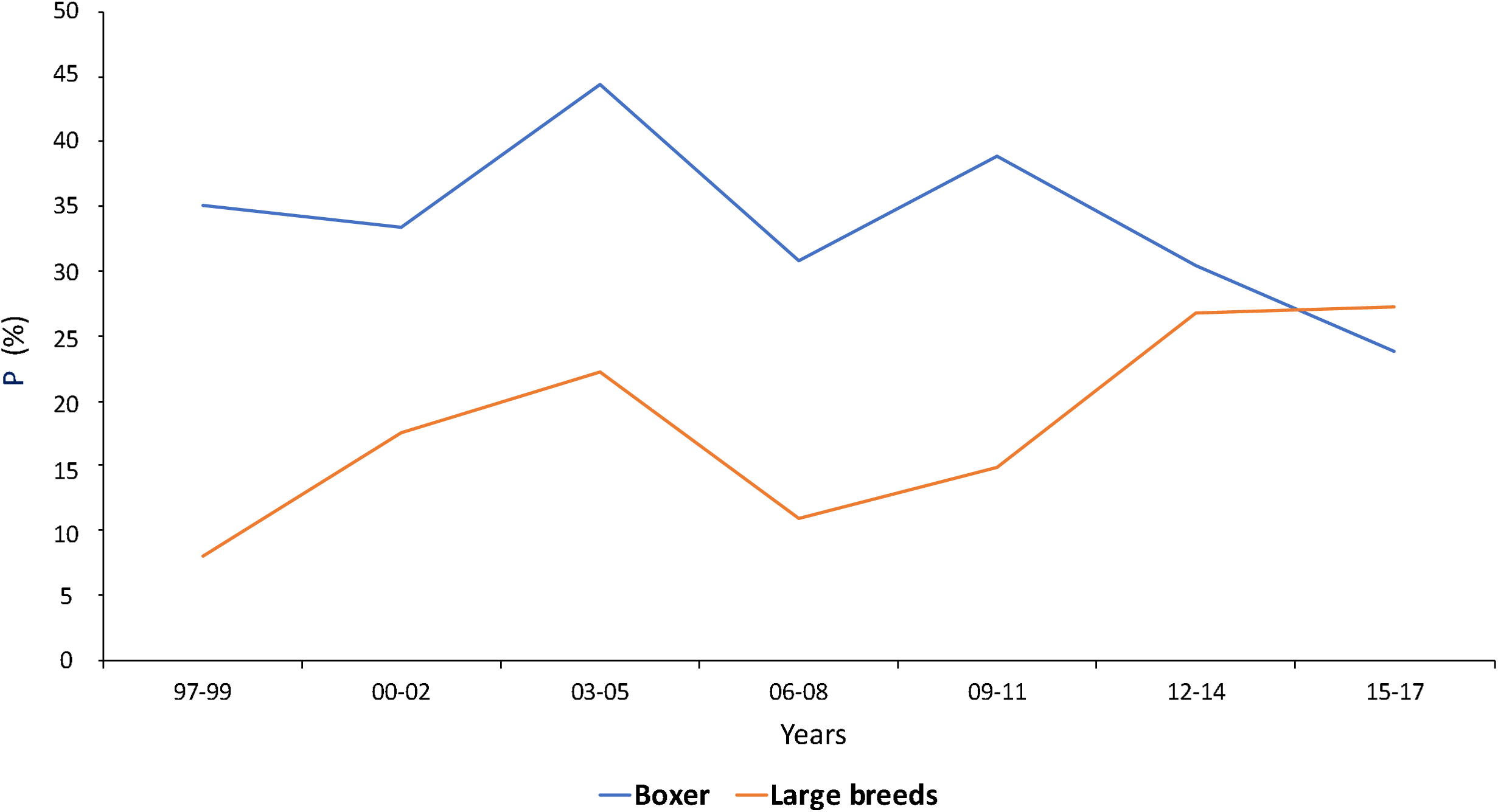
Probability of identifying Pulmonic Stenosis in Boxers and other large breeds (American Staffordshire Terrier; Golden Retriever; German Shepherd; Rottweiler) admitted from 1997 to 2017. Abbreviation: Probability, P.

Since 1963 (The Incredible Journey - Walt Disney) to 2017 (A Dog’s Purpose - Lasse Hallström), Golden Retrievers have been movie stars, which is a well-known reason to explain the increasing popularity of the breed in a social context, and many studies have been performed to explain how media can influence a buyer’s choice [10-12]. PDA was the second most common CHD in our population, in both pure and crossbred dogs. The presence pf PDA was significant in females; therefore, a penetrant autosomal recessive and sex-linked inheritance can be excluded [23]. Although PDA was absent in Boxers, our results indicate the prevalence of PDA was higher than in studies performed in United States and Europe [4-7]. In our study population, PDA was the 2^nd^ most commonly observed CHD; it was frequent in large dog breeds including Dobermanns (89.7%), German Shepherds (65.9%), and Newfoundland (42.9%), as well as in medium and small breeds such as Border Collies (59.1%), Maltese (76.5%), Poodles (65.7%), CKCS (60%) and Chihuahuas (59%). The highest frequency of PDA was observed from 2006 to 2011, and then decreased [5,6,9,33,34]. The reason for the change in the frequency of PDA in that period of time could be explained by the use of Amplatzer Canine Duct Occluder, which is suitable for large breed dogs and that became available in our center in 2006 (Fig 4) [35].

**Figure 4.**
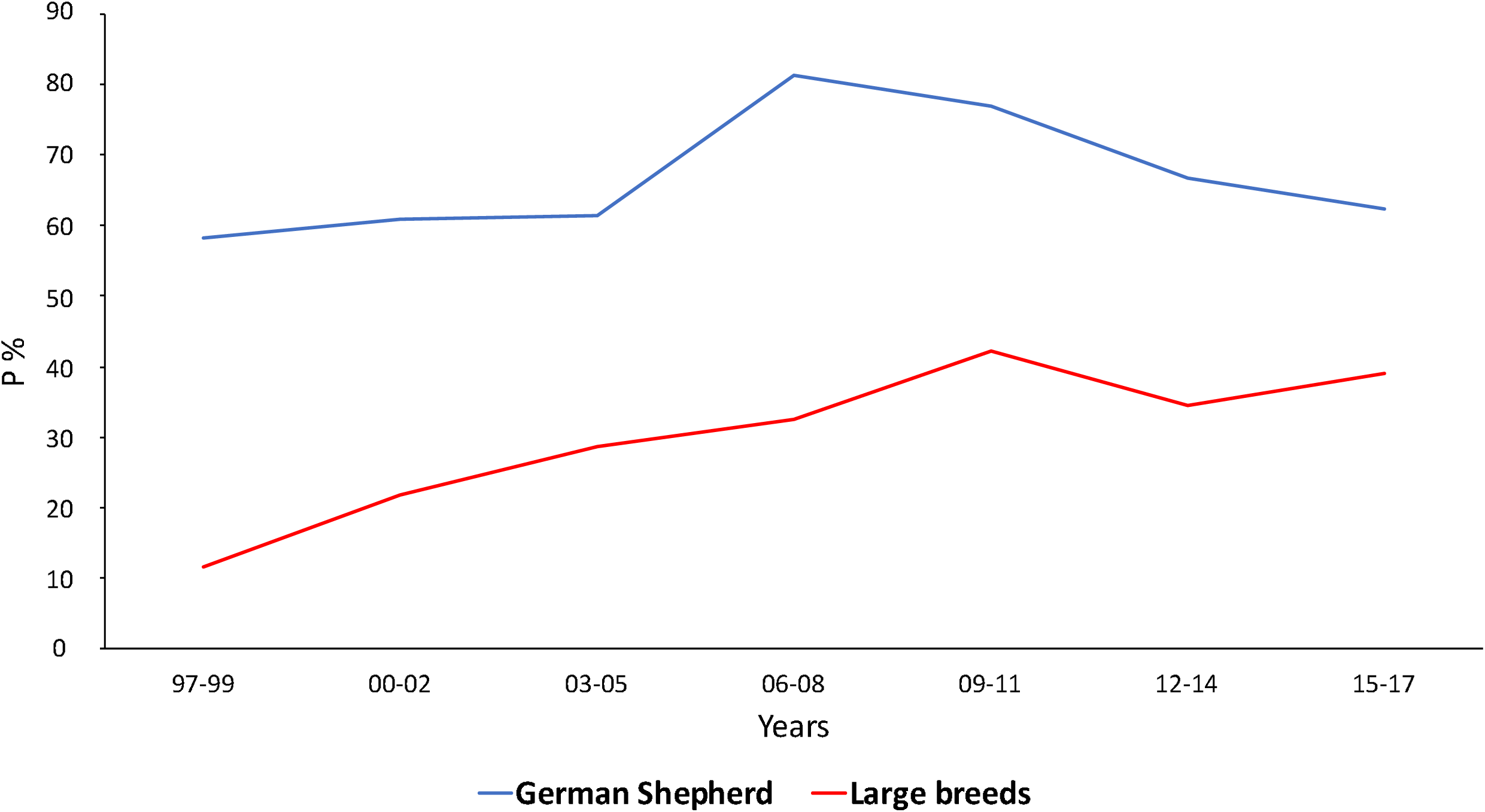
Probability of identifying Patent Ductus Arteriosus in large breeds (Bull Terrier; Dobermann; Golden Retriever; Labrador Retriever; Newfoundland; Rottweiler) admitted from 1997 to 2017. Abbreviation: Probability, P.

Nine cases of rPDA were found in Maltese; it is a very uncommon CHD, and in Maltese has only been described in a publication issued from the same RC in 2011 [9].

SAS was the 3^rd^ most common defect in our study, as a single defect or associated with PS or PDA (8.28%). SAS was found in 72.7% of the Dogue de Bordeaux admitted to the RC from 1997 to 2017, and Type 2 and Type 3 were the most frequent (36.36% each) (Tables 1,2,5).

The three different subtypes were quite equally distributed in Boxers as Type 1 (39.6%), Type 2 (36.63%) and Type 3 (23.76%). It interesting to note that SAS was the 2^nd^ most common CHD in our population from 1997 to 2011, and then its frequency decreased through 2017. SAS and PS are very commonly associated with each other in Boxers (85.8%), and the screening program at this center was aimed to reduce the incidence of both (Fig 4). The reduction of SAS in Boxers is an interesting result because it demonstrates the effectiveness of the screening and breeding program in Boxers. In other words, the increased prevalence of PS in 20 years is not a failure of the Boxer screening and breeding program, but rather the result of the large increase in fashion breeds, such as the French Bulldog, English Bulldog and American Staffordshire Terrier, that are not screened.

AS was the 4^th^ most common CHD in our population. AS was significantly more frequent in males (7.51% CI 4.57– 7.05, P< 0.005) (Table 3) than in females, and Bull Terriers were the most affected breed (30%) (Table 2).

The dogs diagnosed with AS were older than the dogs affected by other CHDs, and the extreme ages at presentation were 50 months (AS) and less than 12 months (TOF) (Fig 1). This result is not surprising because defects of greater severity are associated with the worst symptoms at an early age. Indeed, in many cases, AS is mild in young dogs and becomes progressively worse with age. The murmur in AS can be very soft and necessitate Doppler echocardiographic examination for definitive diagnosis, which is a very different clinical scenario from TOF [5,36].

Many complex defects were found in our population, and PS was the most common CHD detected in association with the other CHDs (SAS, VSD, PDA) (Table 5). The overall prevalence of the PS-SAS association in Boxers (85.8%) seems very high; however, the value has been estimated over the 20-year period (Fig 5). SAS-PDA was very common in Newfoundland; this complex CHDs was found in the 84.6% of the admitted dogs belonging to this breed (Fig 5).

**Figure 5.**
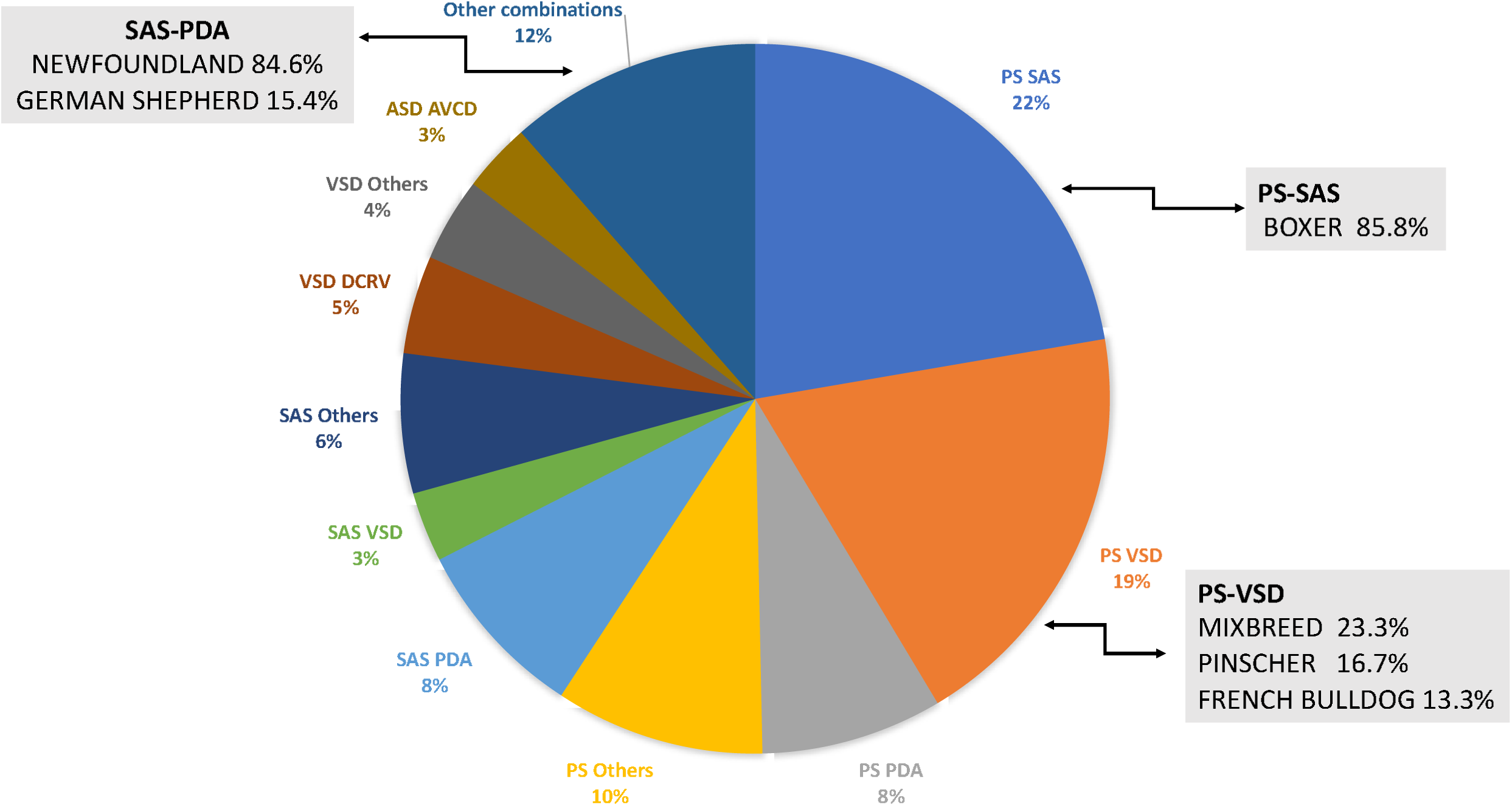
Associated congenital heart defects admitted from 1997 to 2017. Abbreviation: PS, pulmonic stenosis; SAS, subaortic stenosis; VSD, ventricular septal defect; PDA, patent ductus arteriosus, DCRV, double chamber right ventricle; ASD, atrial septal defect; AVCD, atrioventricular canal disease.

PDA and SAS are also very common as simple defects in Newfoundland (Table 2), and the detection of one CHD should be cause for investigation of the other CHDs, in order to exclude the presence of both. The left ventricle volume overload due to a large PDA could cause the overestimation of the severity of SAS. In this case, the correction of PDA determines the reduction of the volume overload, and because the gradient across the aortic valve decreases significantly, the actual severity of subaortic stenosis can be evaluated only after the ductal closure.

Knowing the association among simple CHDs and the breeds involved could be a useful diagnostic tool that should be taken into account in clinical practice.

From 1997 to 2017, several changes have occurred in the clinical and diagnostic approaches to CHDs. The evolution of diagnostic technology, the changing criteria in the classification of some congenital heart diseases, and the increased attention to the selection of breeds prone to CHDs has modified the epidemiological conditions of CHDs in the study RC.

In the final section of the study, the relationships among the popularity, volatility and number of CHDs in individual breeds were investigated over a 20 year time period.

The analyses of the popularity of the breeds found that the number of CHDs detected in a breed increases with the number of registrants of that breed in the ENCI database. This result can be explained by the response to a growing market demand. In this case, the objective of some breeders is to increase the number of puppies of the breed, and little attention is paid to the gene pool strength, to the selection of the ascendants and to a trustworthy breeding program.

Recent studies indicate that breeds with more inherited disorders have become more popular, not less popular, suggesting that health considerations have been secondary in people’s decision to acquire a specific breed of dog [11,14].

Volatility is the average absolute annual change in ENCI registration of dogs belonging to a breed, and it was found to be independent from some breed features (e.g., longer life, inherited genetic disorders, health problems). Societal influences (fashions and fads) have been described as having a primary effect on the popularity of companion breeds, and the volatility of the breeds is an interesting parameter to measure the change in breed popularity over time. The volatility of the French Bulldog was very high (0.10), and the OR of disease was also high (6.98 CI 5.60 – 8.71). In contrast, the volatility of German Shepherds was very low (−0.03) even though the number of registrations of German Shepard puppies is the highest in 20 years among the breeds in our study population. This observation is in accordance with the results obtained by other authors that found that social influence has been more important than functional traits (e.g., health and trainability) in determining owners’ choice of a breed [11,37].

Breed size is thought to be a very important trait behind the owner’s decision to choose a breed. This observation is well supported by the volatility values, with significantly lower or more negative values for large breed dogs, and significantly more positive values for small or medium breed dogs. Chihuahuas, French Bulldogs and CKCS were the most valued small breeds. Among medium size breeds, the Border Collie and American Staffordshire Terrier showed the highest volatility.

However, size was not the only parameter that influenced the popularity of a breed; Yorkshire Terriers and Maltese showed a very low volatility despite their small size.

The influence of media, including movies, television and radio, on the audience is well known and described^10-12^. Unfortunately, people may choose dog breeds based on this media influence and on the idea that a breed is fashionable or a status symbol. These dog owners may not care about the social context in which it should be introduced or the health problems from which a breed may suffer.

The limitations of this study were primarily associated with its retrospective nature; some cases could not be included because of a lack of clinical and diagnostic information. In particular, the absence of information about the prevalent breeds in crossbred dog has been a limitation in identifying relationships between breed and CHD in this class of dogs. A bias could also arise because the study was conducted in a single cardiological referral center that specializes in the surgical or percutaneous repair of PS, PDA, VSD and ASD, and this center has been unique in our country for a long period of time. This specialization of this particular center is the reason why CHD and some breeds (e.g., Boxer, German Shepherd) are overrepresented in our study population. However, this specialization could also be a point of strength because any variation in the preferences of breeds can be monitored from a consistent study location.

## Conclusions

In conclusion, this study allowed us to evaluate the Boxer screening program for CHDs, whose success is evidenced by the decreased prevalence of SAS and PS in this breed.

However, the paradox that people buy breeds of dog that are predisposed to congenital heart diseases was also evidenced in our study, and, as reported elsewhere, fashions and trends influence many individual choices [11,12,29,30]. The owners are not often fully aware of the potential problems their dog may face prior to acquisition of a dog [31,32]. It is also possible that owners do not perceive the clinical signs of some inherited cardiac disorders as problems, but rather as normal, breed-specific characteristics (e.g., murmur in CKCS).

In general, when choosing a breed, owners may consider other characteristics to be more important than dog health. Nevertheless, the authors think that an effective breeding program should start with educating the owners about the health problems of a breed. If the owners are not motivated to buy a healthy breed, then breeds with inherent health problems will be perpetuated, and the motivation of breeders to address health problems in their breed reduced.

In this context, the importance of creating a network of veterinary cardiology centers that monitor the distribution of a breed and treat the problem of CHDs using the same clinical approach and diagnostic procedures is clear. This approach could be a useful instrument to provide breeders with effective support in implementing the breeding program in order to control the diffusion of CHDs, without impoverishing the genetic pool.

## Acknowledgments

The authors thank Dr.Fabrizio Crivellari (Executive Director), Dr. Dino Muto (President) and Dr. Selene Festa, for providing the Italian Kennel Club (ENCI) dataset essential in this study.

